# The effects of antibiotic combination treatments on *Pseudomonas aeruginosa* tolerance evolution and coexistence with *Stenotrophomonas maltophilia*

**DOI:** 10.1101/2022.03.25.485776

**Authors:** Jack P. Law, A. Jamie Wood, Ville-Petri Friman

## Abstract

*Pseudomonas aeruginosa* bacterium is a common pathogen of Cystic Fibrosis (CF) patients due to its ability to evolve resistance to antibiotics during treatments. While *P. aeruginosa* resistance evolution is well characterised in monocultures, it is less well understood in polymicrobial CF infections. Here, we investigated how exposure to ciprofloxacin, colistin, or tobramycin antibiotics, administered at sub-MIC doses alone and in combination, shaped the tolerance evolution of *P. aeruginosa* (PAO1 lab and clinical CF LESB58 strains) in the absence and presence of a commonly co-occurring species, *Stenotrophomonas maltophilia*. Increases in antibiotic tolerances were primarily driven by the presence of that antibiotic in the treatment. We observed a reciprocal cross-tolerance between ciprofloxacin and tobramycin, and when combined these antibiotics selected increased MICs for all antibiotics. Though the presence of *S. maltophilia* did not affect the tolerance or the MIC evolution, it drove *P. aeruginosa* into extinction more frequently in the presence of tobramycin due to its relatively greater innate tobramycin tolerance. In contrast, *P. aeruginosa* dominated and drove *S. maltophilia* extinct in most other treatments. Together, our findings suggest that besides driving high-level antibiotic tolerance evolution, sub-MIC antibiotic exposure can alter competitive bacterial interactions, leading to target pathogen extinctions in multi-species communities.

## Introduction

Cystic Fibrosis (CF) is a genetic condition that is characterised by impaired chloride ion channel function, resulting in thick mucus secretions in the lungs that are susceptible to chronic bacterial infection (1). Of the bacterial species that infect adult CF patients, *Pseudomonas aeruginosa* is the most prevalent pathogen associated with morbidity (2, 3) and is difficult to treat due to its intrinsic resistance to many antibiotics and its ability to readily evolve resistance to new antibiotics (4, 5). Though usually dominated by *P. aeruginosa*, CF infections are often polymicrobial and many different bacterial species co-occur with *P. aeruginosa* in the CF lungs (6–10).

Over the course of their lives patients with CF will be treated with a number of different antibiotics; for example, during treatment to eradicate *P. aeruginosa* or to help resolve pulmonary exacerbations (11, 12). Antibiotics are administered at high concentration multiple times per day to maintain a therapeutic dose at a concentration greater than the minimum inhibitory concentration (MIC) required to inhibit bacterial growth (12–14). Antibiotic combinations are used to target multiple species simultaneously or to increase efficacy against a single species (15, 16). However, the thick mucus secretions and complex branching structure of the lungs themselves will likely result in bacterial populations experiencing a gradient of antibiotic concentrations (17, 18). Thus, despite the best efforts of the treatment regimens, pockets of bacteria within the lungs are likely to experience antibiotic concentrations below that required to inhibit those bacteria, and such subinhibitory concentrations have been shown to promote antibiotic resistance evolution (19, 20).

Selection for antibiotic resistance differs between antibiotics administered at-or-above MIC, and below MIC (14). At concentrations greater than MIC the driver of selection is whether the bacteria can survive the antibiotic challenge, and so any mutations in the bacterial population that increase MIC, regardless of the impact on other competitive growth traits, would be selected (19, 20). Conversely, below MIC the selective pressure differs, such that any mutation that confers an increase in growth in the presence of the low antibiotic concentration, and thus a competitive advantage, relative to other members of the population would be selected regardless of whether this mutation would increase the MIC (19, 20). This relatively weaker selection pressure increases the number of viable mutations, which in turn increases the likelihood that one such mutation could increase MIC through mechanisms not traditionally considered to be involved in resistance (20). The lack of antibiotic-mediated killing also results in a longer selective window during which more mutations can accumulate and either ameliorate the costs of higher-level resistance (21), or together confer high-level resistance via epistatic interactions (22). While effects of lethal concentrations (>MIC) of antibiotic combinations on individual bacterial species have been explored previously (23, 24), they are less well understood at sub-MIC concentrations in multi-species communities.

Competition with other bacterial species could change the trajectory of antibiotic resistance evolution in a focal pathogen species in various ways (25). Firstly, the presence of competitors more tolerant of an antibiotic treatment than the susceptible pathogen species could increase the strength of competition between the two and lead to a decrease in relative pathogen abundance, potentially triggering extinctions (26). Competitor-mediated reduction in the population density of the focal pathogen could further slow resistance evolution by reducing the mutation supply rate and the emergence of *de novo* resistance (27). When antibiotic resistance evolves it is often associated with metabolic costs, such as activation of efflux pumps or modification of the antibiotic target. These costs could reduce pathogen competitiveness in the presence of non-resistant mutants, or species unaffected by a given antibiotic, when antibiotic concentrations are low (28–30). While it has been suggested that bacterial interactions are predominantly competitive (31), it is also possible that other interacting bacteria could facilitate antibiotic resistance of the focal species via horizontal gene transfer of resistance genes (32). Alternatively, other species could provide protection from antibiotics via secretions that break down antibiotics (21, 33, 34) or create protective microenvironments via production of biofilms (30, 35). Despite being a ubiquitous selective force in nature, there are relatively few studies directly testing the effect of bacterial inter-species interactions on the evolution of antibiotic resistance.

Here we focused on studying how the evolution of antibiotic tolerance of *P. aeruginosa* is affected by the presence of *Stenotrophomonas maltophilia*, another CF-associated species that is increasing in prevalence among CF patients (36–41) and commonly co-occurs with *P. aeruginosa* (42). In order to investigate this, we performed a short-term *in vitro* serial transfer experiment in which we grew both of the lung-naïve laboratory *P. aeruginosa* strain PAO1 and the lung-adapted Liverpool Epidemic Strain B58 (LESB58; 42) either alone in monoculture or in the presence of *S. maltophilia* (resulting in four different cultures: PAO1, LESB58, PAO1 & *S. maltophilia*, and LESB58 & *S. maltophilia*). Two strains of *P. aeruginosa* were chosen to compare the potential effect of previous exposure to antibiotic treatments and other infecting bacteria on the evolution of antibiotic tolerance. Each of these cultures were treated with one of the eight combinations (see Methods) of the anti-Pseudomonal antibiotics ciprofloxacin, colistin, and tobramycin. These antibiotics were selected because of their use in either *P. aeruginosa* eradication therapy or for treatment of pulmonary exacerbations (11, 12, 44, 45), and for their differing modes of action (46–48). Each of the antibiotics were applied at a sub-MIC concentration that had small but contrasting effects on the growth of all three bacterial strains (Supplementary Figure 1).

During the serial transfer experiment, which took place over 21 days, we tracked the presence of *P. aeruginosa* and *S. maltophilia* for any extinctions and monitored changes in total population densities across the 192 selection lines. Following the experiment, we measured the ability of the evolved focal *P. aeruginosa* isolates to grow in the treatment concentrations of the individual antibiotics relative to ancestral stock strains. Moreover, the MICs of each antibiotic were determined for all evolved *P. aeruginosa* isolates. We hypothesised that: i) antibiotic tolerance evolution could be constrained in the presence of a competitor but promoted in the presence of multiple antibiotics if cross-tolerance evolution is common; and ii) antibiotic exposure could change community composition due to differences between the species’ innate susceptibility to the antibiotics or due to evolution of tolerance-growth trade-offs.

We found increases in antibiotic tolerance or MIC were not generally enhanced by antibiotic combinations; rather increases in tolerance or MIC to a given antibiotic were driven by the presence of that antibiotic in the treatment combination, which occasionally led to cross-tolerance. Similarly, the presence of *S. maltophilia* did not affect antibiotic tolerance evolution or MIC with either *P. aeruginosa* strain. However, while both *P. aeruginosa* strains were able to dominate the “No Antibiotic” coculture treatments, tobramycin-containing antibiotic treatments triggered *P. aeruginosa* extinctions in 15% of coculture populations, which were more common when *S. maltophilia* was cultured with PAO1 than LESB58. Together, these results suggest that the effects of sub-MIC antibiotic concentrations could be magnified in polymicrobial communities due to competition and asymmetry in innate antibiotic tolerances.

## Materials and Methods

### Bacterial strains and culture conditions

Two strains of *Pseudomonas aeruginosa* were used as the focal pathogen species: PAO1, a lab adapted reference strain (ATCC 15692), and LESB58, a transmissible CF lung isolated strain (42). *Stenotrophomonas maltophilia* type strain ATCC 13637—isolated from the oropharyngeal tract of a cancer patient (49)—was used as the coculture competitor. The base media used throughout was a 50:50 mix of nutrient broth without NaCl (Sigma; 5 g/l peptic digest of animal tissue, 3 g/l beef extract, pH 6.9) and PBS (8 g/l NaCl, 2 g/l KCl, 1.42 g/l Na_2_HPO_4_, 2.4 g/l KH_2_PO_4_), hereafter ‘NB’, that allowed stable coexistence of both the focal pathogen (*P. aeruginosa*) and competitor (*S. maltophilia*) species over a single 72 h growth period. All cultures, unless otherwise stated, were grown at 37°C with shaking at 180 rpm.

### Selection experiment

During the selection experiment, a focal bacterium (either *P. aeruginosa* strain PAO1 or LESB58) was grown in a culture either alone (monoculture) or with *S. maltophilia* (coculture) and treated with subinhibitory concentrations of ciprofloxacin (CIP), colistin (CST), and tobramycin (TOB) antibiotics in all one-, two- and three-way combinations (“No Antibiotic”, CIP, CST, TOB, CIP+CST, CIP+TOB, CST+TOB, CIP+CST+TOB). Each treatment was replicated 6 times for both focal pathogens in the absence and presence of *S. maltophilia* resulting in a total of 192 selection lines. During the initial setup overnight cultures, from frozen stocks, of PAO1, LESB58, and *S. maltophilia*, were diluted down to the same optical density at 600 nm (OD_600_; approximately 0.17 at 600 nm), corresponding to cell densities of 7.4×10^6^, 2.2×10^7^, and 4.6×10^6^ CFU/ml respectively. Monocultures consisted of 20 μl of the *P. aeruginosa* strain, while cocultures mixed 10 μl of the *P. aeruginosa* strain with 10 μl of *S. maltophilia*, each in 200 μl of NB supplemented with one of the eight antibiotic treatments for a total volume of 220 μl. The concentrations of the antibiotics (0.03215 μg/ml ciprofloxacin (Sigma Aldrich), 2 μg/ml colistin (Acros Organics), and 0.5 μg/ml tobramycin (Acros Organics)) were chosen to be below the minimum inhibitory concentration of all three species (as determined below; Table 1, Supplementary Figure 1), and were kept constant across each combination. Selection lines were grown in 96-well plates. To measure the dynamics of the bacterial densities, all replicates were homogenised by mixing and OD_600_ of each replicate was measured every 72 hours (Tecan Infinite 200). After measurement, each replicate was again mixed and 20 μl of the culture was transferred to 200 μl of fresh media with the same antibiotic treatment, which were incubated for 72 hours until the next serial transfer. Presence or absence of each species in each culture was determined following each transfer by growing subsamples of the 72-hour cultures on different selective agar; *Pseudomonas* selective agar (Oxoid; *Pseudomonas* agar base: 16 g/l gelatin peptone, 10 g/l casein hydrolysate, 10 g/l potassium sulphate, 1.4 g/l magnesium chloride, 11 g/l agar, 1% vol/vol glycerol; *Pseudomonas* CN selective supplement: 200 μg/ml centrimide, 15 μg/ml sodium nalidixate), and *S. maltophilia* selective agar: LB agar (10 g/l tryptone, 5 g/l yeast extract, 5 g/l NaCl, 15 g/l agar) supplemented with 64 μg/ml tobramycin incubated at 30°C rather than 37°C— *S. maltophilia* is innately resistant towards tobramycin at 30°C (50). Some monoculture replicates were contaminated with *S. maltophilia* (14 PAO1 selection lines, and 9 LESB58 selection lines), and these were excluded from the analyses. Whole population bacterial samples were picked from the agar plates for each replicate, grown overnight in NB, and cryopreserved in 20% glycerol to be frozen at −80°C. The selection experiment was carried out for 21 days, equalling 6 serial transfers.

**Table 1:** Minimum inhibitory concentrations for the three ancestral bacterial strains, along with the experimental concentrations used of each antibiotic.

| Antibiotic | Treatment concentration (µg/ml) | Minimum inhibitory concentration (µg/ml)<br>[treatment concentration as proportion of MIC] |  |  |
| --- | --- | --- | --- | --- |
|  |  | PAO1 | LESB58 | *S. maltophilia* |
| Ciprofloxacin | 0.03125 | 0.0625 | 1 | 0.125 |
|  |  | [1/2] | [1/32] | [1/4] |
| Colistin | 2 | 4 | 4 | 8 |
|  |  | [1/2] | [1/2] | [1/4] |
| Tobramycin | 0.5 | 1 | 2 | 8 |
|  |  | [1/2] | [1/4] | [1/16] |

### Determination of MIC and antibiotic tolerance

Both prior to and following the selection experiment, the minimum inhibitory concentration (MIC) of each of the three antibiotics—ciprofloxacin, colistin, and tobramycin—was determined by broth microdilution for the three bacteria strains. Briefly, overnight cultures, from frozen stocks, were diluted 1 in 10 in PBS and further diluted 1 in 10 into NB with antibiotic concentrations ranging from 32 μg/ml to 0.015625 μg/ml (2^5^ to 2^-6^) and grown in static conditions, in triplicate. OD_600_ was measured after 24 hours (Tecan Sunrise). MIC was defined as the lowest concentration of antibiotic at which there was no growth. For the evolved strains, the MIC_50_ of a bacterial population was defined as the MIC required to inhibit half of the replicates of that population.

To assess the difference in tolerance of the antibiotics, at the treatment concentrations used during the selection experiments, for each individual replicate we define a growth proxy, 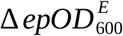, as

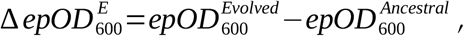

where 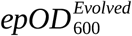 is the endpoint OD_600_ from growth after 24 hours of the evolved strain in a given antibiotic treatment, and analogously for the ancestral strain. Such a measure is a proxy for the growth of the bacteria and we use it accordingly in Figure 1 and its associated statistics.

**Figure 1:**
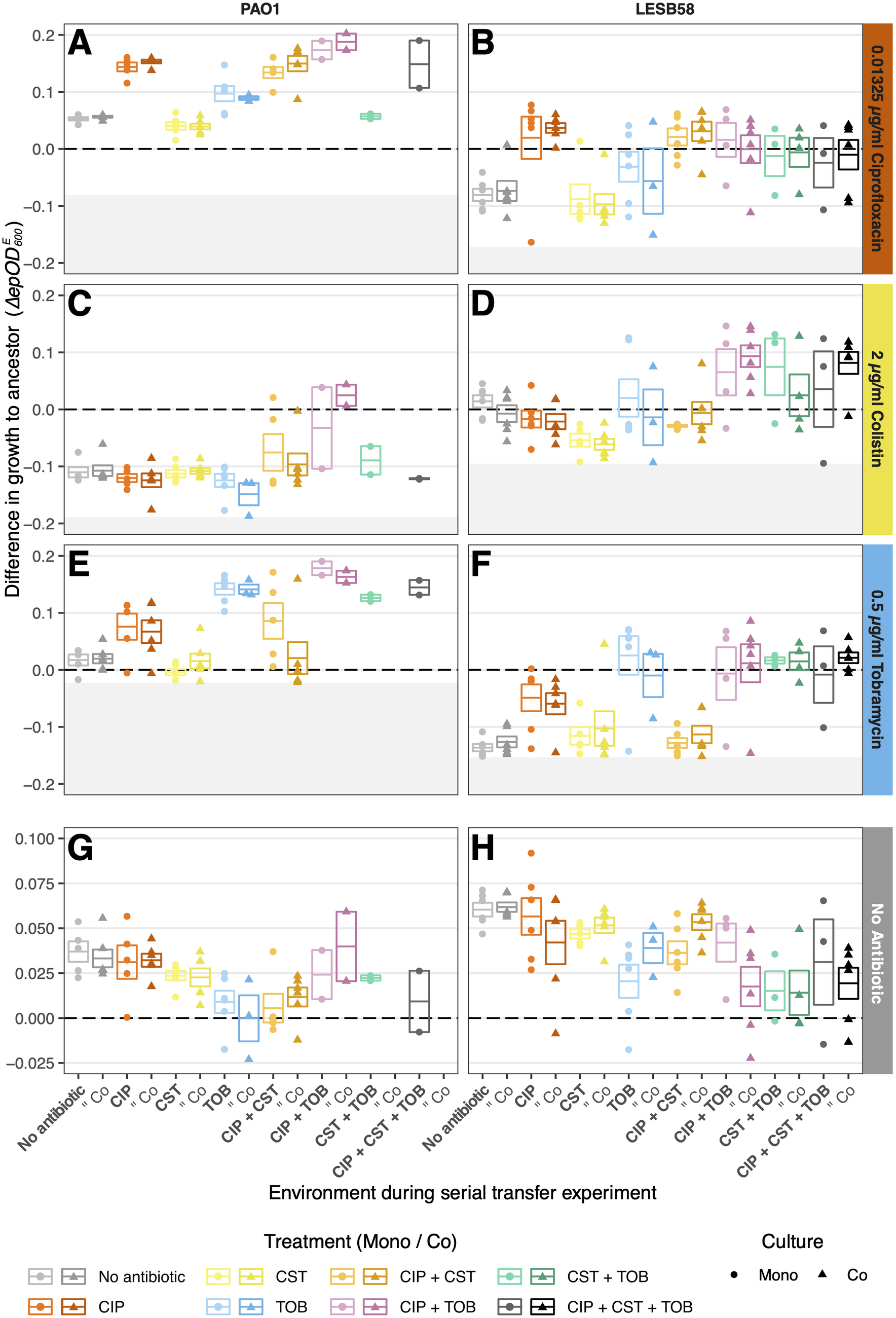
Growth of each evolved *P. aeruginosa* replicate population, relative to their respective ancestor, in the same treatment concentrations of each antibiotic (A–F) or without antibiotic (G–H), measured in separate growth assays at the end of the selection experiment. Panel columns show the two *P. aeruginosa* strain, while panel rows show growth in the presence and absence of different antibiotics. The horizontal dashed line represents growth equal to that of the ancestor (i.e., relative change in OD_600_ = 0). Each point represents the mean growth for three technical replicates of one replicate population, minus the growth of a similarly replicated ancestral growth (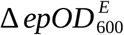, as defined in Methods). Boxes show mean of all populations (centre line; 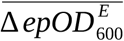, as defined in Methods), and upper and lower limits represent ±S.E.M. Shapes show mono- (circle; ‘Mono’) and cocultures (triangle; ‘Co’); colours show antibiotic treatment, with lighter and darker shades representing the absence and presence of the *S. maltophilia* competitor. Shaded grey portions of the panels represents no growth, i.e., 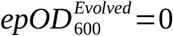. Cross tolerance between ciprofloxacin and tobramycin can be seen in panels A, B and E, F, comparing “No Antibiotic” treatment in columns 1, 2, to CIP and TOB treatments in columns 3, 4 and 7, 8. Cost of tobramycin tolerance can be seen in panels G, H, comparing “No Antibiotic” in columns 1, 2, to TOB treatments in columns 7, 8.

We use an aggregate measure when considering the difference in tolerance to ancestral strains of evolved bacteria from a selection regimen as a whole

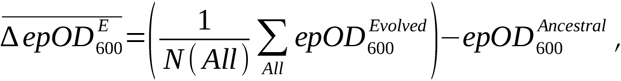

where we calculate the errors in our measure by computing the standard error of the mean (S.E.M.) of the evolved strains only. The sum is over all relevant strains evolved in the selection regimen of interest, e.g., monoculture PAO1 treated with CIP, and N ≤ 6 replicates dependent on extinctions. All bacteria (ancestral and evolved) were also grown without antibiotic in NB for 24 hours, and difference in growth without antibiotic was calculated as above.

### Statistical analyses

All data were analysed in R version 3.6.3 (51). Data manipulation and graphing were performed using the *tidyverse* suite of packages (52), along with *egg* for figure assembly (53), and *ggbeeswarm* for point plotting (54). Regarding the tolerance dataset, separate linear regression models for each *P. aeruginosa* species were used when analysing each response variable (i.e., growth difference relative to the ancestor during CIP exposure, CST exposure, TOB exposure, and “No Antibiotic” exposure). Here, the response variable was difference in growth relative to the ancestor 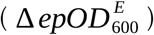, as defined above. Two-way type II ANOVA was performed using the *car* package (55). Post-hoc pairwise comparisons were computed from estimated marginal means using Tukey’s honest significance test, through the `emmeans` and contrast functions of the *emmeans* package (56). Pairwise comparisons were computed between treatments alone, after observing no effect of competitor nor interaction. Full pairwise comparisons can be found in Supplementary Spreadsheet 1.

Regarding the MIC dataset, individual Pearson Chi-squared tests of independence were performed for each *P. aeruginosa* strain in each antibiotic. MIC values were represented as ordered nominal variables, and frequency of observed MIC for each replicate in each treatment was tabulated. Chi-squared tests were computed using the chisq_test function from the *coin* package (57). Pairwise tests of independence with Benjamini-Hochberg false discovery rate corrections were performed between each treatment using the `pairwiseOrdinalIndependence` function from the *rcompanion* package (58). Full pairwise comparisons can be found in Supplementary Spreadsheet 2.

With the population density dataset, two linear regression models were fit, one to each *P. aeruginosa* strain. The response variable was natural logarithm transformed OD_600_ values, and antibiotic treatment and competitor as the predictor variables. Twoway type II ANOVA and post-hoc pairwise comparisons were performed as with the tolerance data. Full pairwise comparisons can be found in Supplementary Spreadsheet 3.

## Results

### Effects of antibiotic treatments on *P. aeruginosa* antibiotic tolerance and relative cost of tolerance

To test potential tolerance evolution, we compared the ancestral and evolved *P. aeruginosa* populations’ ability to grow in the presence of the treatment concentration of each of the antibiotics. Some *P. aeruginosa* monoculture replicates were removed from the analyses due to contamination with *S. maltophilia* (see Methods). Moreover, as *P. aeruginosa* went extinct in some of the tobramycin-containing treatments (24/143 selection lines), the evolution of tolerance was compared using only surviving treatment replicates.

Firstly, the antibiotic tolerances of both strains were not affected by previous exposure to *S. maltophilia* (Figure 1, *p* > 0.05, Supplementary Table 1). With regards to the control treatments, in the case of the clinical isolate LESB58 the “No Antibiotic” control treatment resulted in increased susceptibility to antibiotics relative to the ancestor, while the antibiotic treatments maintained the ancestral-level tolerances of ciprofloxacin and tobramycin. In contrast, the “No Antibiotic” control treatment of the lab strain PAO1 maintained ancestral-level tolerance, while the antibiotic treatments further increased the tolerance of the evolved isolates. More specifically, with both strains there was a significant increase in ciprofloxacin tolerance when treated with the ciprofloxacin (CIP) mono-, CIP+colistin (CST) and CIP+tobramycin (TOB) treatments compared to the “No Antibiotic” control treatment (Pairwise Tukey post-hoc, PAO1: *t*(48) = 10.46 (CIP); 9.72 (CIP+CST); 10.27 (CIP+TOB), *p* < 0.001; LESB58: *t*(66) = 4.57 (CIP); 4.47 (CIP+CST); 3.47 (CIP+TOB), *p* < 0.001; Figure 1A, B). Similarly, the TOB and CIP+TOB treatments significantly increased tobramycin tolerance in PAO1 (Pairwise Tukey post-hoc, *t*(48) = 6.48 (TOB); 6.38 (CIP + TOB), *p* < 0.001; Figure 1E), while all tobramycin containing treatments significantly increased tolerance in LESB58 (Pairwise Tukey post-hoc, *t*(66) = 5.61 (TOB); 5.69 (CIP+TOB); 5.66 (CST+TOB); 5.56 (CIP+CST+TOB), *p* < 0.001; Figure 1F). In contrast, colistin-containing treatments did not generally result in increased colistin tolerance: for both strains no colistin-containing treatments resulted in any statistical change in colistin tolerance compared to the “No Antibiotic” control treatments (Pairwise Tukey post-hoc, *p* > 0.05; Figure 1C, D).

We also observed cross-tolerance between ciprofloxacin and tobramycin, i.e., the CIP mono-treatment provided tobramycin tolerance, and vice versa (Figure 1). In PAO1, the CIP mono-treatment trended towards increased tobramycin tolerance (Pairwise Tukey post-hoc, *t*(48) = 3.03, *p* = 0.07; Figure 1E), while TOB monotreatment gave a significant increase in ciprofloxacin tolerance compared to the “No Antibiotic” control treatment (Pairwise Tukey post-hoc, *t*(48) = 3.94, *p* = 0.006). However, the increase in ciprofloxacin cross-tolerance from the TOB mono-treatment was not as great as resulted from the CIP mono-treatment (Pairwise Tukey post-hoc, *t*(48) = 5.69, *p* < 0.001; Figure 1A). A similar pattern emerged in LESB58, where the CIP mono-treatment resulted in a significantly higher tobramycin tolerance than the “No Antibiotic” control (Pairwise Tukey post-hoc, *t*(66) = 5.61, *p* < 0.001; Figure 1F), and, though not significant, a majority of replicates treated with TOB or CST+TOB had a greater ciprofloxacin tolerance than the mean “No Antibiotic” control (Figure 1B). Additionally, the CIP+TOB combination treatment resulted in cross-tolerance to colistin in both strains (Pairwise Tukey post-hoc, PAO1: *t*(48) = 5.01, *p* < 0.001; LESB58: *t*(66) = 3.32, *p* = 0.03; Figure 1B, F). Colistin-containing treatments did not provide any cross-tolerance towards the other antibiotics, and in the case of tobramycin tolerance the combination of CIP+CST did not lead to cross-tolerance as in the CIP mono-treatment (Figure 1C, G). These results suggest that while colistin tolerance evolution was rare, both pathogen strains readily evolved tolerance to ciprofloxacin and tobramycin, which was driven by prior exposure to these antibiotics and reciprocal cross-tolerance.

To test whether selection in different antibiotic treatments led to a cost of tolerance, we grew each of the surviving evolved replicates in media without antibiotic and compared their growth relative to the respective ancestors (Figure 1G, H). The majority of both *P. aeruginosa* genotype replicates across all treatments evolved to grow better in the growth media relative to their ancestors (Figure 1G, H), and the increase in growth, relative to the ancestor, was greater in the lung-adapted LESB58 than the lab-adapted PAO1. However, this increase clearly varied between the antibiotic treatments. In the case of both genotypes, the TOB mono-treatment constrained adaptation, resulting in a significantly reduced growth compared to the “No Antibiotic” control treatment (Pairwise Tukey post-hoc, PAO1: *t*(44) = 4.47, *p* = 0.001; LESB58: *t*(66) = 3.40, *p* = 0.02). Moreover, the growth of evolved LESB58 populations treated with any tobramycin-containing antibiotic treatment were significantly below the “No Antibiotic” control treatment (Pairwise Tukey post-hoc, *t*(66) = 3.58, *p* = 0.01 (CIP+TOB); 4.81, *p* < 0.001 (CST+TOB); 3.89, *p* = 0.005 (CIP+CST+TOB); Figure 1H). These results suggest that adapting to tolerate tobramycin reduced the growth and potential competitive ability of *P. aeruginosa* strains.

### Changes in the MIC of antibiotics with evolved *P. aeruginosa* populations

We measured changes in the Minimum Inhibitory Concentration (MIC) of each antibiotic for evolved *P. aeruginosa* replicate populations, and MIC_50_—the MIC capable of inhibiting 50% of replicates—for each treatment to quantify whether exposure to low antibiotic concentrations led to increased MIC. For both *P. aeruginosa* strains across all three antibiotics, there was no effect of previous exposure to *S. maltophilia* on the MICs. However, the MICs of the evolved populations changed considerably with both *P. aeruginosa* strains in response to all antibiotics (Supplementary Table 2).

#### Changes in ciprofloxacin MIC

Both evolved *P. aeruginosa* strains showed large increases in MIC to ciprofloxacin (Figure 2A, B). The MIC of ciprofloxacin for PAO1 replicates from the “No Antibiotic” control treatment remained mostly unchanged relative to the ancestor, at 0.125 μg/ml, though a pair of individual replicates increased their MIC by three-fold (Figure 2A); whereas in LESB58 the baseline effect of the “No Antibiotic” control was a three-fold decrease in MIC compared to the ancestor, from 1 μg/ml to 0.125 μg/ml—to the same MIC as the laboratory PAO1 strain (Figure 2B). Pairwise Chi-squared tests showed that the CIP, CIP+CST, and CIP+TOB treatments all resulted in significantly greater MIC values, compared to the control treatment, amongst isolates of both strains (Pairwise independence; PAO1—CIP: *X*^2^(1, N = 22) = 15.61, *p* = 0.002; CIP+CST: *X*^2^(1, N = 22) = 10.77, *p* = 0.010; CIP+TOB: *X*^2^(1, N = 15) = 8.34, *p* = 0.018; Figure 2A; LESB58—CIP: *X*^2^(1, N = 24) = 19.03, *p* > 0.001; CIP+CST: *X*^2^(1, N = 24) = 13.71, *p* = 0.0019; CIP+TOB: *X*^2^(1, N = 22) = 13.31, *p* = 0.0019; Figure 2B). Indeed, in LESB58 the triple antibiotic treatment also significantly increased MIC values (Pairwise independence: *X*^2^(1, N = 21) = 9.95, *p* = 0.0064; Figure 2B), such that all ciprofloxacin-containing treatments increased ciprofloxacin MIC. Furthermore, many of the TOB mono-treated isolates from both strains had high MIC values, and in LESB58 the MICs for both these and the CST+TOB treated isolates were significantly different to the “No Antibiotic” control treatment (Pairwise independence; TOB: *X*^2^(1, N = 21) = 7.34, *p* = 0.019; CST+TOB: *X*^2^(1, N = 19) = 10.03, *p* = 0.0064), further suggesting that there is some cross-tolerance provided by tobramycin as also seen in the growth measurements (Figure 1). The MIC values for evolved LESB58 isolates reached higher levels than in PAO1, with 18 LESB58 isolates reaching 4 or 8 μg/ml compared with one PAO1 isolate, and there was also greater variation in MIC values among the LESB58 isolates of a given treatment than PAO1. Overall, both *P. aeruginosa* strains evolved an increase in ciprofloxacin MIC, which was driven mostly by the previous exposure to ciprofloxacin.

**Figure 2:**
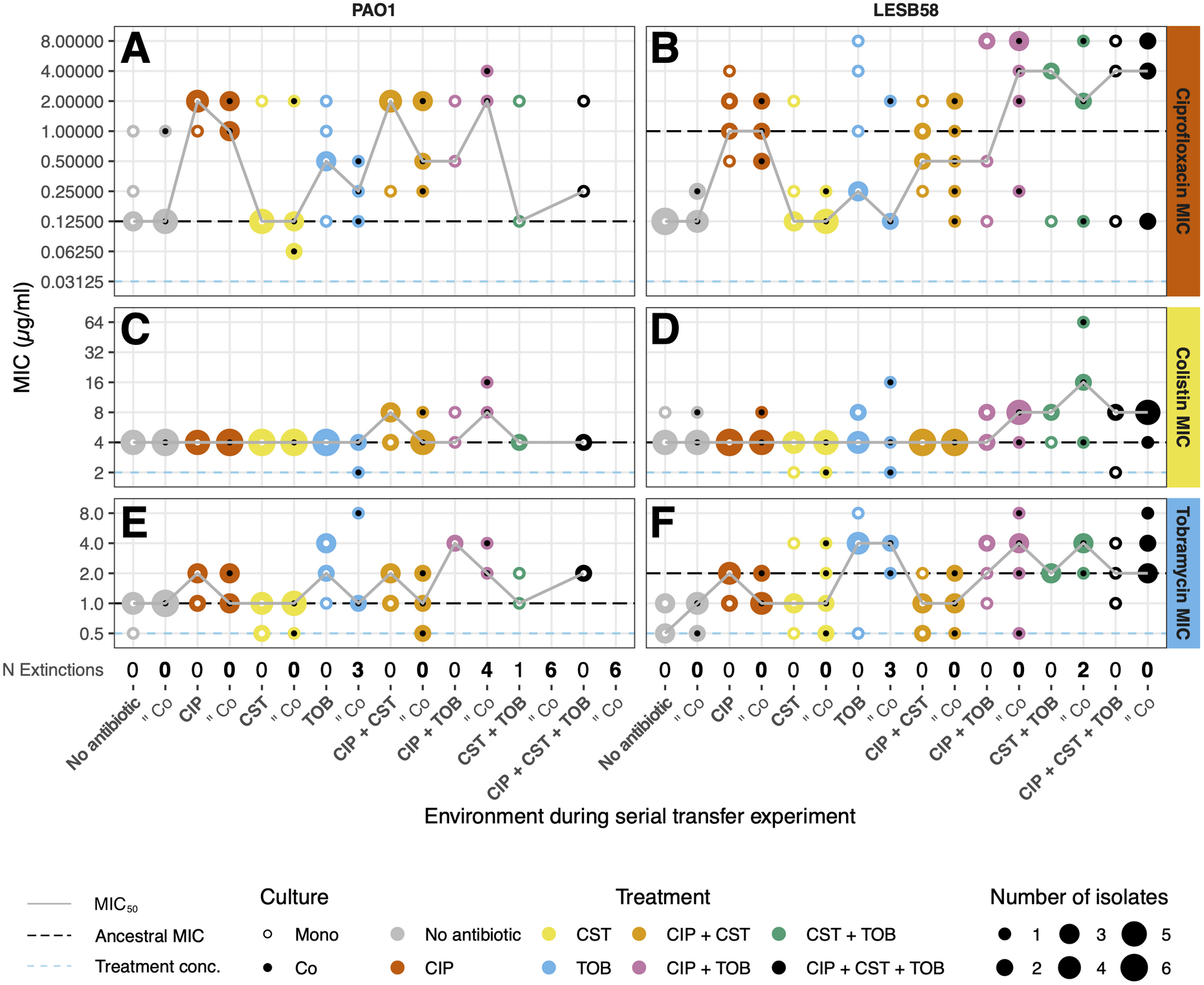
The MIC of the three individual antibiotics for each evolved replicate population of *P. aeruginosa* strains. Panel columns show *P. aeruginosa* strain and rows the MIC of each antibiotic. The dashed line represents the MIC of the respective ancestors; the grey line shows the MIC_50_ of each treatment across replicate populations, as defined in the methods; the dashed blue line shows the treatment concentration. The size of each point represents the number of replicates with the specified MIC. Colour of the points represents treatment; white centre dot represents monoculture and black represents coculture. Number of extinctions in each treatment and coculture shown beneath the X-axis. MIC was measured in triplicate for each replicate. Cross tolerance between ciprofloxacin and tobramycin can be seen in panels A, B and E, F, comparing “No Antibiotic” treatment in columns 1, 2, to CIP and TOB treatments in columns 3, 4 and 7, 8.

#### Changes in colistin MIC

As opposed to ciprofloxacin, the MICs of colistin did not increase as a result of prior colistin exposure during the selection experiment (Figure 2C, D). In the case of PAO1, the majority of treatments resulted in no change to the ancestral MIC of 4 μg/ml (Figure 2C). Slightly more variation was observed amongst the LESB58 isolates that had been exposed to CIP, CST, or TOB mono-treatments, though no changes greater than one-fold for more than a single replicate were found. However, the MICs of isolates treated with the combinations CIP+TOB and CST+TOB, were both significantly higher compared with the “No Antibiotic” control treatment (Pairwise independence; CIP+TOB: X2(1, N = 22) = 6.13, p = 0.037; CST+TOB: X2(1, N = 19) = 6.21, p = 0.037; Figure 2C). Overall, only small changes in colistin MIC were observed, which were indirectly driven by other antibiotics.

#### Changes in tobramycin MIC

The MIC changes for tobramycin were similar between both *P. aeruginosa* strains (Figure 2E, F). The “No Antibiotic” control treated isolates of PAO1 maintained the ancestral MIC of 1 μg/ml (Figure 2E), whereas the “No Antibiotic” control treated LESB58 isolates decreased in MIC relative to their ancestor (from 2 μg/ml down to 0.5 and 1 μg/ml; Figure 2F). For both strains, the TOB and CIP+TOB treatments resulted in significant increase in MIC compared with the “No Antibiotic” control treatment (Pairwise independence; PAO1—TOB: *X*^2^(1, N = 20) = 8.54, *p* = 0.0035; CIP+TOB: *X*^2^(1, N = 15) = 12.00, *p* < 0.001; Figure 2E; LESB58—TOB: *X*^2^(1, N = 21) = 12.67, *p* = 0.0035; CIP+TOB: *X*^2^(1, N = 22) = 11.35, *p* = 0.0053; Figure 2F), and for LESB58 this was the also the case with the CST+TOB and triple antibiotic treatments (Pairwise independence; CST+TOB: *X*^2^(1, N = 19) = 13.77, *p* = 0.0035; CIP+CST+TOB: *X*^2^(1, N = 21) = 13.22, *p* = 0.0035). Both strains also had a significant increase in MIC compared to the “No Antibiotic” control treatment as a result of the CIP monotreatment (Pairwise independence; PAO1: *X*^2^(1, N = 22) = 7.98, *p* = 0.019); LESB58: *X*^2^(1, N = 24) = 10.58, *p* = 0.0064), though this increase did not reach the same values as the tobramycin containing treatments did: the MIC of the CIP mono-treated LESB58 isolates was significantly lower compared to TOB, CST+TOB, and CIP+CST+TOB treatments (Pairwise independence; TOB: *X*^2^(1, N = 21) = 7, *p* = 0.021; CST+TOB:*X*^2^(1, N = 19) = 8.05, *p* = 0.014; CIP+CST+TOB: *X*^2^(1, N = 21) = 6.64, *p* = 0.023), suggesting that the cross-tolerance provided by ciprofloxacin was weaker than that of tobramycin. Overall, both *P. aeruginosa* strains evolved increases in tobramycin MIC, which was primarily driven by the previous exposure to tobramycin during the selection experiment.

### The effect of antibiotic treatments on bacterial densities and coculture composition

We measured a proxy of total population density of each bacterial population at the final timepoint of the selection experiment to determine the extent to which antibiotics inhibited bacterial growth (measurements were taken at the population level and did not differentiate species frequencies; optical density at 600 nm). For both *Pseudomonas* strains, there was a significant effect of antibiotic treatment on total population density (Supplementary Table 3, *p* < 0.001), and with the exception of the CST mono-treatment, antibiotics generally decreased the total population density relative to the “No Antibiotic” control treatment (Figure 3A–D). However, post-hoc pairwise comparisons showed that this effect was driven by the cocultures, as none of the monocultures differed significantly between any of the treatments with either strain (Figure 3A, B). The combination of CST+TOB was particularly effective in the cocultures of both strains, reducing the population significantly compared with the “No Antibiotic” control (Pairwise Tukey post-hoc, PAO1: *t*(65) = 4.21, *p* = 0.0041; LESB58: *t*(71) = 3.98, *p* = 0.0081; Figure 3C, D). There was no effect of growing in monoculture vs. coculture for PAO1 (Supplementary Table 3, *p* > 0.05), and though there was a significant effect in LESB58 (Supplementary Table 3, *p* = 0.022) this was likely driven by the large difference in population density between the two CST-mono treated cultures (Pairwise Tukey post-hoc, *t*(71) = 3.62, *p* = 0.024; Figure 3B, D). Together, these results show that the antibiotics reduced bacterial population densities compared with the “No Antibiotic” control treatment regardless of the presence of *S. maltophilia*, and that the combination of CST+TOB was highly effective at reducing total bacterial population densities.

**Figure 3:**
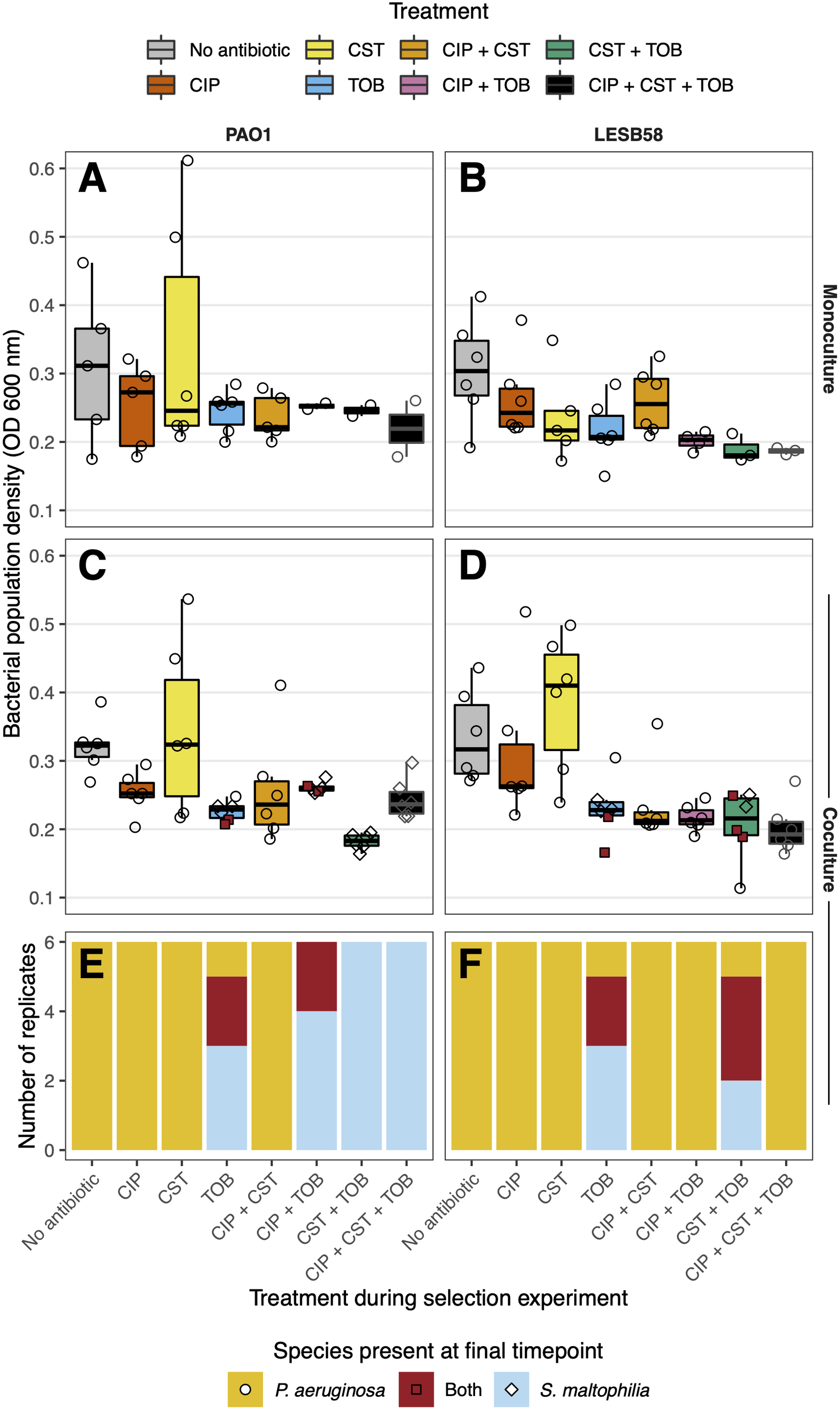
Total bacterial population densities and composition of cocultures at the final timepoint of the selection experiment. **A–D:** Boxplots of total bacterial population densities (optical density at 600 nm) of each replicate population from eight treatments (see legend for boxplot fill colours). Panel columns show the *P. aeruginosa* strain, rows monocultures or cocultures. Points represent individual replicates; N = 6. Shapes show species present at the final timepoint: *P. aeruginosa* as circles, *S. maltophilia* as diamonds, and both as red squares. **E–F:** The presence of surviving species in each coculture replicate (N = 6). Colours represent the surviving species as follows: *P. aeruginosa* in orange, *S. maltophilia* in blue, and both in red.

We also compared the composition of *P. aeruginosa* and *S. maltophilia* cocultures at the end of the experiment to examine the effects of antibiotics on the species coexistence. We found that *P. aeruginosa* survived in all monocultures across the different treatments (except for a single PAO1 replicate that went extinct under CST+TOB treatment). This suggests that the low concentrations of the antibiotic were not sufficient to kill *P. aeruginosa*, even when applied in combination. However, extinctions of *P. aeruginosa* were more common in the presence of *S. maltophilia* (Figure 3E, F). Interestingly, in the absence of antibiotics, both *P. aeruginosa* genotypes were able to dominate the cocultures, driving *S. maltophilia* extinct in all replicates. While the same was true in the CIP and CST treatments, the TOB mono-treatment allowed prolonged coexistence between the two bacteria and *S. maltophilia* was able to survive with both *P. aeruginosa* genotypes in two of the six TOB mono-treatment replicates, and fully outcompeted *P. aeruginosa* in a further three replicates (Figure 3E, F). Furthermore, the two *P. aeruginosa* genotypes differed in their capacity to coexist with *S. maltophilia* across the antibiotic combination treatments, with the laboratory strain PAO1 driven extinct more often than the clinical strain LESB58; PAO1 was only able to survive in two CIP+TOB treated replicates and was otherwise driven extinct in the remaining 22 replicates across the other tobramycin-containing combination treatments (Figure 3E). In contrast, the LESB58 strain dominated *S. maltophilia* in all combination treatments except for the CST+TOB treatment, in which *P. aeruginosa* was driven extinct in two replicates (Figure 3F). Together, these results suggest that while *P. aeruginosa* was able to outcompete *S. maltophilia* in most of the environments, this relationship was reversed in the presence of tobramycin, leading to either *P. aeruginosa* extinction or coexistence with *S. maltophilia*.

## Discussion

While antibiotics are routinely used to treat *P. aeruginosa* infections within the polymicrobial communities in CF patient lungs, it is unclear how low antibiotic concentrations affect antibiotic resistance evolution in the presence of naturally occurring CF lung microbiota. Here we studied this by exposing *P. aeruginosa* to sub-MIC concentrations of ciprofloxacin, colistin, tobramycin and their combinations in the presence and absence of a commonly co-occurring CF species, *S. maltophilia*. We observed *P. aeruginosa* tolerance evolution to all antibiotics and a clear cross-tolerance between tobramycin and ciprofloxacin. Moreover, antibiotic exposure often led to an increase in MIC, suggesting that sub-MIC selection can lead to high levels of antibiotic tolerance. While the presence of a competitor had no effect on antibiotic tolerance evolution, antibiotic exposure had a strong effect on species community composition: even though *P. aeruginosa* dominated most of the treatments, it coexisted with-, or was driven into extinction by, *S. maltophilia* in the presence of tobramycin due to innate differences in tobramycin resistance. Thus, even low doses of antibiotics could significantly change antibiotic tolerance evolution and composition of multi-species communities.

We predicted that antibiotic tolerance evolution could be constrained by the presence of *S. maltophilia* as a competitor, for example due to negative effects on *P. aeruginosa* population densities and mutation supply rate (27). However, we found that the presence of *S. maltophilia* did not alter the trajectory of antibiotic tolerance evolution. Instead, *P. aeruginosa* evolved increased tolerance to all antibiotics regardless of the competitor presence, which was driven primarily by previous exposure to the same antibiotics during the selection experiment. We also predicted that presence of multiple antibiotics could select for increased levels of antibiotic tolerance potentially due to cross-tolerance or via selection for generalised resistance mechanisms such as efflux pumps (4, 5). In support of this we found that increasing the numbers of antibiotics in the treatments resulted in higher level MICs compared to mono-antibiotic treatments. We also found a reciprocal cross-tolerance between ciprofloxacin and tobramycin in both *P. aeruginosa* strains.

While ciprofloxacin and tobramycin resistance can be mediated by the same mechanism in *P. aeruginosa*—upregulation of the MexXY-OprM efflux pump (59, 60)—previous studies have suggested that *in vitro* selection for such mutations are rare (61, 62). As a result, other antibiotic-specific resistance mechanisms could have evolved, such as mutations in *fusA1* for tobramycin (62–64) and *gyrAB* for ciprofloxacin (29, 65), even though these mutations are not known to provide crosstolerance to the other antibiotic. Interestingly, in LESB58 the CST+TOB combination resulted in high levels of MICs for all three antibiotics, providing a cross tolerance to ciprofloxacin. Colistin and tobramycin resistance can be mediated by outer membrane modifications via activation of the PmrAB (66–68) and ParRS (69, 70) two component systems. Gain of function mutations in either *pmrB* or *parS* can result in increased tolerance to both antibiotics and have been observed in *P. aeruginosa* treated with aminoglycosides *in vitro* (62, 71) and in the clinic (22, 68). However, whether the decrease in membrane permeability that these systems provide is sufficient to prevent entry of ciprofloxacin is unclear. We also found that sub-MIC antibiotic selection often led to a clear increase in MIC, providing tolerance to much higher concentrations of antibiotics than the bacteria experienced during the selection experiment. With PAO1, this was especially clear with the ciprofloxacin and tobramycin MICs, while LESB58 showed increase in MIC to all antibiotics but mainly when exposed to antibiotic combinations. It has been shown previously that low levels of antibiotic selection can lead to high levels of resistance due to epistasis (20), or that antibiotic resistance can evolve *de novo* even in the absence of antibiotic selection due to adaptation to the growth media (72). Further genetic analyses are required to ascertain the genetic mechanisms of antibiotic resistance in sub-MIC concentrations and to better understand the molecular basis of cross-tolerance.

We also predicted that antibiotic exposure could alter the community composition via potential differences in innate antibiotic sensitivity or tolerance-growth trade-offs. The baseline interaction between *P. aeruginosa* and *S. maltophilia* in the “No Antibiotic” control cocultures was antagonistic, whereby *P. aeruginosa* competitively excluded *S. maltophilia*. Indeed, it has previously been shown that *P. aeruginosa* can kill *S. maltophilia* via a contact dependent mechanism during planktonic growth, and that *P. aeruginosa* outcompetes *S. maltophilia* when grown in dual-species biofilms (73, 74). However, while competitive exclusion of *S. maltophilia* was observed in the absence and presence of most antibiotic treatments, this pattern was reversed in the presence of tobramycin where coexistence with the innately resistant *S. maltophilia* (50) or extinction of *P. aeruginosa* was observed. Though no follow-up data exist for the extinct *P. aeruginosa* replicates in the tobramycin-containing cocultures, it is possible that they were unable to evolve tolerance to tobramycin, compared to surviving replicates, and were thus competitively excluded from cocultures. Moreover, evolution of tolerance by *P. aeruginosa* did not restore competitive dominance, as surviving coculture replicates that evolved tobramycin tolerance most frequently coexisted with *S. maltophilia*. The most probable explanation for this was the cost of tobramycin tolerance, which resulted in reduced growth of evolved *P. aeruginosa* isolates and thus likely less intense competition between the two species, though other mechanisms cannot be ruled out.

Competitive exclusion of *P. aeruginosa* differed between the two strains: the laboratory strain PAO1 was driven extinct in each tobramycin combination except for two CIP+TOB replicates, whereas the clinical strain LESB58 was only affected in the CST+TOB treatment. This could, to some degree, be explained by relatively higher ciprofloxacin and tobramcyin tolerance of LESB58, which could have reduced the negative effect of antibiotics in the CIP+TOB and triple antibiotic treatments compared to PAO1. The initial differences in antibiotic susceptibility could reflect contrasting evolutionary histories between these two strains. PAO1 is highly lab-adapted due to repeated culturing in lab media while LESB58, a transmissible strain isolated in 1988 from CF patients (43), is adapted to the CF lungs by producing more biofilm than PAO1 (75) and lacks motility (42). Importantly, LESB58 has a greater interbacterial competitive ability—producing greater amounts of competitive factors, such as pyocyanin and proteases, and secreting these earlier in its growth phases (76, 77)— than PAO1 which is likely beneficial during polymicrobial CF infections when in competition with other bacteria (78, 79). LESB58 was thus likely pre-adapted to compete with other species such as *S. maltophilia*, which could explain the lower frequency of extinctions. Antibiotic treatments containing tobramycin were more effective at driving competitive exclusion in the naïve PAO1 strain, suggesting that during early infection—when failure to eradicate *P. aeruginosa* can select for mutations that aid establishment of long-term chronic infections (80)—combination antibiotic treatments may be more effective at clearing *P. aeruginosa* than individual treatments. Of the antibiotic combinations, colistin and tobramycin together resulted in extinction of both strains, adding support for their efficacy in clinic (15). Together, our findings suggest that the efficacy of antibiotic combinations can be magnified in polymicrobial infections, leading to higher clearance of the target pathogen.

In summary, while the presence of competitor did not affect the trajectory of antibiotic tolerance evolution in *P. aeruginosa*, we found that exposure to sublethal antibiotic concentrations led to more frequent extinctions of the target pathogen in the presence of an antibiotic resistant competitor. Specifically, tobramycin played a key role in this ecological process and its effects persisted even after evolution of tobramycin tolerance by *P. aeruginosa* likely due to the associated growth costs, leading to coexistence and weakening of *P. aeruginosa* dominance. Somewhat worryingly, *P. aeruginosa* populations often evolved increases in MIC to all antibiotics, leading to resistance against much higher concentrations of antibiotics than experienced during the selection experiment. In conclusion, our results suggest that differences in antibiotic susceptibility can magnify competition in bacterial communities, leading to changes in community composition. The efficiency of the antibiotic treatment is then determined by both the surrounding community as well as efficacy of delivery, choice of antibiotic, and antibiotic concentration, further complicating treatment design.

## Author Contributions

All authors conceived and designed the study, JPL collected and analysed the data, and all authors wrote the manuscript.

## References

1. Hart CA, Winstanley C. 2002. Persistent and aggressive bacteria in the lungs of cystic fibrosis children. Br Med Bull 61:81–96.

2. Orenti A, Zolin A, Jung A, van Rens J. 2021. European Cystic Fibrosis Society Patient Registry Annual Data Report 2019. URL: https://www.ecfs.eu/projects/ecfs-patient-registry/annual-reports

3. UK Cystic Fibrosis Registry. 2021. UK Cystic Fibrosis Registry Annual data report 2020. URL: https://www.cysticfibrosis.org.uk/the-work-we-do/uk-cf-registry/reporting-and-resources

4. Poole K. 2011. Pseudomonas Aeruginosa: Resistance to the Max. Front Microbiol 2:1–13.

5. López-Causapé C, Cabot G, del Barrio-Tofiño E, Oliver A. 2018. The Versatile Mutational Resistome of Pseudomonas aeruginosa. Front Microbiol 9:1–9.

6. Zhao J, Schloss PD, Kalikin LM, Carmody LA, Foster BK, Petrosino JF, Cavalcoli JD, VanDevanter DR, Murray S, Li JZ, Young VB, LiPuma JJ. 2012. Decade-long bacterial community dynamics in cystic fibrosis airways. Proc Natl Acad Sci 109:5809–5814.

7. Kramer R, Sauer-Heilborn A, Welte T, Jauregui R, Brettar I, Guzman CA, Höfle MG. 2015. High Individuality of Respiratory Bacterial Communities in a Large Cohort of Adult Cystic Fibrosis Patients under Continuous Antibiotic Treatment. PLoS One 10:e0117436.

8. Coburn B, Wang PW, Diaz Caballero J, Clark ST, Brahma V, Donaldson S, Zhang Y, Surendra A, Gong Y, Elizabeth Tullis D, Yau YCW, Waters VJ, Hwang DM, Guttman DS. 2015. Lung microbiota across age and disease stage in cystic fibrosis. Sci Rep 5:10241.

9. Zemanick ET, Wagner BD, Robertson CE, Ahrens RC, Chmiel JF, Clancy JP, Gibson RL, Harris WT, Kurland G, Laguna TA, McColley SA, McCoy K, Retsch-Bogart G, Sobush KT, Zeitlin PL, Stevens MJ, Accurso FJ, Sagel SD, Harris JK. 2017. Airway microbiota across age and disease spectrum in cystic fibrosis. Eur Respir J 50:1700832.

10. Einarsson GG, Zhao J, LiPuma JJ, Downey DG, Tunney MM, Elborn JS. 2019. Community analysis and co-occurrence patterns in airway microbial communities during health and disease. ERJ Open Res 5:00128–02017.

11. UK Cystic Fibrosis Trust. 2009. Antibiotic Treatment for Cystic Fibrosis. Third Edition. London. URL: https://www.cysticfibrosis.org.uk/the-work-we-do/resources-for-cf-professionals/consensus-documents

12. Langton Hewer SC, Smyth AR. 2017. Antibiotic strategies for eradicating Pseudomonas aeruginosa in people with cystic fibrosis. Cochrane Database Syst Rev 2020.

13. Heijerman H, Westerman E, Conway S, Touw D. 2009. Inhaled medication and inhalation devices for lung disease in patients with cystic fibrosis: A European consensus. J Cyst Fibros 8:295–315.

14. Andersson DI, Hughes D. 2012. Evolution of antibiotic resistance at non-lethal drug concentrations. Drug Resist Updat 15:162–172.

15. Herrmann G, Yang L, Wu H, Song Z, Wang H, Høiby N, Ulrich M, Molin S, Riethmüller J, Döring G. 2010. Colistin□Tobramycin Combinations Are Superior to Monotherapy Concerning the Killing of Biofilm Pseudomonas aeruginosa. J Infect Dis 202:1585–1592.

16. Barbosa C, Beardmore R, Schulenburg H, Jansen G. 2018. Antibiotic combination efficacy (ACE) networks for a Pseudomonas aeruginosa model. PLOS Biol 16:e2004356.

17. Moriarty TF, McElnay JC, Elborn JS, Tunney MM. 2007. Sputum antibiotic concentrations: Implications for treatment of cystic fibrosis lung infection. Pediatr Pulmonol 42:1008–1017.

18. Bos AC, Passé KM, Mouton JW, Janssens HM, Tiddens HAWM. 2017. The fate of inhaled antibiotics after deposition in cystic fibrosis: How to get drug to the bug? J Cyst Fibros 16:13–23.

19. Gullberg E, Cao S, Berg OG, Ilbäck C, Sandegren L, Hughes D, Andersson DI. 2011. Selection of Resistant Bacteria at Very Low Antibiotic Concentrations. PLoS Pathog 7:e1002158.

20. Wistrand-Yuen E, Knopp M, Hjort K, Koskiniemi S, Berg OG, Andersson DI. 2018. Evolution of high-level resistance during low-level antibiotic exposure. Nat Commun 9:1599.

21. Bottery MJ, Wood AJ, Brockhurst MA. 2017. Adaptive modulation of antibiotic resistance through intragenomic coevolution. Nat Ecol Evol 1:1364–1369.

22. Jochumsen N, Marvig RL, Damkiær S, Jensen RL, Paulander W, Molin S, Jelsbak L, Folkesson A. 2016. The evolution of antimicrobial peptide resistance in Pseudomonas aeruginosa is shaped by strong epistatic interactions. Nat Commun 7:13002.

23. Vestergaard M, Paulander W, Marvig RL, Clasen J, Jochumsen N, Molin S, Jelsbak L, Ingmer H, Folkesson A. 2016. Antibiotic combination therapy can select for broad-spectrum multidrug resistance in Pseudomonas aeruginosa. Int J Antimicrob Agents 47:48–55.

24. Barbosa C, Mahrt N, Bunk J, Graßer M, Rosenstiel P, Jansen G, Schulenburg H. 2021. The Genomic Basis of Rapid Adaptation to Antibiotic Combination Therapy in Pseudomonas aeruginosa. Mol Biol Evol 38:449–464.

25. Letten AD, Hall AR, Levine JM. 2021. Using ecological coexistence theory to understand antibiotic resistance and microbial competition. Nat Ecol Evol 5:431–441.

26. Bottery MJ, Pitchford JW, Friman V-P. 2021. Ecology and evolution of antimicrobial resistance in bacterial communities. ISME J 15:939–948.

27. de Visser JAGM, Rozen DE. 2006. Clonal Interference and the Periodic Selection of New Beneficial Mutations in Escherichia coli. Genetics 172:2093–2100.

28. Stickland HG, Davenport PW, Lilley KS, Griffin JL, Welch M. 2010. Mutation of nfxB Causes Global Changes in the Physiology and Metabolism of Pseudomonas aeruginosa. J Proteome Res 9:2957–2967.

29. Jørgensen KM, Wassermann T, Jensen PØ, Hengzuang W, Molin S, Høiby N, Ciofu O. 2013. Sublethal Ciprofloxacin Treatment Leads to Rapid Development of High-Level Ciprofloxacin Resistance during Long-Term Experimental Evolution of Pseudomonas aeruginosa. Antimicrob Agents Chemother 57:4215–4221.

30. Klümper U, Recker M, Zhang L, Yin X, Zhang T, Buckling A, Gaze WH. 2019. Selection for antimicrobial resistance is reduced when embedded in a natural microbial community. ISME J 13:2927–2937.

31. Foster KR, Bell T. 2012. Competition, Not Cooperation, Dominates Interactions among Culturable Microbial Species. Curr Biol 22:1845–1850.

32. Botelho J, Grosso F, Peixe L. 2019. Antibiotic resistance in Pseudomonas aeruginosa – Mechanisms, epidemiology and evolution. Drug Resist Updat 44:100640.

33. Bottery MJ, Wood AJ, Brockhurst MA. 2016. Selective Conditions for a Multidrug Resistance Plasmid Depend on the Sociality of Antibiotic Resistance. Antimicrob Agents Chemother 60:2524–2527.

34. Bottery MJ, Matthews JL, Wood AJ, Johansen HK, Pitchford JW, Friman V-P. 2022. Inter-species interactions alter antibiotic efficacy in bacterial communities. ISME J 16:812–821.

35. Beaudoin T, Yau YCW, Stapleton PJ, Gong Y, Wang PW, Guttman DS, Waters V. 2017. Staphylococcus aureus interaction with Pseudomonas aeruginosa biofilm enhances tobramycin resistance. npj Biofilms Microbiomes 3:25.

36. Talmaciu I, Varlotta L, Mortensen J, Schidlow D V. 2000. Risk factors for emergence ofStenotrophomonas maltophilia in cystic fibrosis. Pediatr Pulmonol 30:10–15.

37. Marchac V, Equi A, Le Bihan-Benjamin C, Hodson M, Bush A. 2004. Casecontrol study of Stenotrophomonas maltophilia acquisition in cystic fibrosis patients. Eur Respir J 23:98–102.

38. Spicuzza L, Sciuto C, Vitaliti G, Dio G, Leonardi S, Rosa M. 2009. Emerging pathogens in cystic fibrosis: ten years of follow-up in a cohort of patients. Eur J Clin Microbiol Infect Dis 28:191–195.

39. Salsgiver EL, Fink AK, Knapp EA, LiPuma JJ, Olivier KN, Marshall BC, Saiman L. 2016. Changing Epidemiology of the Respiratory Bacteriology of Patients With Cystic Fibrosis. Chest 149:390–400.

40. Hatziagorou E, Orenti A, Drevinek P, Kashirskaya N, Mei-Zahav M, De Boeck K, Pfleger A, Sciensano MT, Lammertyn E, Macek M, Olesen HV, Farge A, Naehrlich L, Ujhelyi R, Fletcher G, Padoan R, Timpare Z, Malakauskas K, Fustik S, Gulmans V, Turcu O, Pereira L, Mosescu S, Rodic M, Kayserova H, Krivec U, Vazquez-Cordero C, de Monestrol I, Lindblad A, Jung A, Makukh H, Carr SB, Cosgriff R, Zolin A. 2020. Changing epidemiology of the respiratory bacteriology of patients with cystic fibrosis–data from the European cystic fibrosis society patient registry. J Cyst Fibros 19:376–383.

41. Capaldo C, Beauruelle C, Saliou P, Rault G, Ramel S, Héry-Arnaud G. 2020. Investigation of Stenotrophomonas maltophilia epidemiology in a French cystic fibrosis center. Respir Med Res 78:100757.

42. Dalbøge CS, Hansen CR, Pressler T, Høiby N, Johansen HK. 2011. Chronic pulmonary infection with Stenotrophomonas maltophilia and lung function in patients with cystic fibrosis. J Cyst Fibros 10:318–325.

43. Winstanley C, Langille MGI, Fothergill JL, Kukavica-Ibrulj I, Paradis-Bleau C, Sanschagrin F, Thomson NR, Winsor GL, Quail MA, Lennard N, Bignell A, Clarke L, Seeger K, Saunders D, Harris D, Parkhill J, Hancock REW, Brinkman FSL, Levesque RC. 2009. Newly introduced genomic prophage islands are critical determinants of in vivo competitiveness in the Liverpool Epidemic Strain of Pseudomonas aeruginosa. Genome Res 19:12–23.

44. Mogayzel PJ, Naureckas ET, Robinson KA, Mueller G, Hadjiliadis D, Hoag JB, Lubsch L, Hazle L, Sabadosa K, Marshall B. 2013. Cystic Fibrosis Pulmonary Guidelines. Am J Respir Crit Care Med 187:680–689.

45. Smith S, Rowbotham NJ, Regan KH. 2018. Inhaled anti-pseudomonal antibiotics for long-term therapy in cystic fibrosis. Cochrane Database Syst Rev 2018.

46. Kotra LP, Haddad J, Mobashery S. 2000. Aminoglycosides: Perspectives on Mechanisms of Action and Resistance and Strategies to Counter Resistance. Antimicrob Agents Chemother 44:3249–3256.

47. Bialvaei AZ, Samadi Kafil H. 2015. Colistin, mechanisms and prevalence of resistance. Curr Med Res Opin 31:707–721.

48. Hooper DC, Jacoby GA. 2016. Topoisomerase Inhibitors: Fluoroquinolone Mechanisms of Action and Resistance. Cold Spring Harb Perspect Med 6:a025320.

49. Hugh R, Leifson E. 1963. A description of the type strain of Pseudomonas maltophilia. Int Bull Bacteriol Nomencl Taxon 13:133–138.

50. Mooney L, Kerr K., Denton M. 2001. Survival of Stenotrophomonas maltophilia following exposure to concentrations of tobramycin used in aerosolized therapy for cystic fibrosis patients. Int J Antimicrob Agents 17:63–66.

51. R Core Team. 2019. R: A Language and Environment for Statistical Computing. Vienna, Austria.

52. Wickham H. 2017. tidyverse: Easily Install and Load the “Tidyverse.”

53. Auguie B. 2019. egg: Extensions for “ggplot2”: Custom Geom, Custom Themes, Plot Alignment, Labelled Panels, Symmetric Scales, and Fixed Panel Size.

54. Clarke E, Sherrill-Mix S. 2017. ggbeeswarm: Categorical Scatter (Violin Point) Plots.

55. Fox J, Weisberg S, Price B. 2019. car: Companion to Applied Regression.

56. Lenth R. 2019. emmeans: Estimated Marginal Means, aka Least-Squares Means.

57. Hothorn T, Hornik K, Wiel MA van de, Zeileis A. 2008. Implementing a Class of Permutation Tests: The coin Package. J Stat Softw 28:1–23.

58. Mangiafico S. 2020. rcompanion: Functions to Support Extension Education Program Evaluation.

59. Masuda N, Sakagawa E, Ohya S, Gotoh N, Tsujimoto H, Nishino T. 2000. Substrate Specificities of MexAB-OprM, MexCD-OprJ, and MexXY-OprM Efflux Pumps in Pseudomonas aeruginosa. Antimicrob Agents Chemother 44:3322–3327.

60. Hocquet D, Vogne C, El Garch F, Vejux A, Gotoh N, Lee A, Lomovskaya O, Plésiat P. 2003. MexXY-OprM Efflux Pump Is Necessary for Adaptive Resistance of Pseudomonas aeruginosa to Aminoglycosides. Antimicrob Agents Chemother 47:1371–1375.

61. Yen P, Papin JA. 2017. History of antibiotic adaptation influences microbial evolutionary dynamics during subsequent treatment. PLOS Biol 15:e2001586.

62. López-Causapé C, Rubio R, Cabot G, Oliver A. 2018. Evolution of the Pseudomonas aeruginosa Aminoglycoside Mutational Resistome In Vitro and in the Cystic Fibrosis Setting. Antimicrob Agents Chemother 62:1–6.

63. Scribner MR, Santos-Lopez A, Marshall CW, Deitrick C, Cooper VS. 2020. Parallel Evolution of Tobramycin Resistance across Species and Environments. MBio 11:1–17.

64. Bolard A, Plésiat P, Jeannot K. 2018. Mutations in Gene fusA1 as a Novel Mechanism of Aminoglycoside Resistance in Clinical Strains of Pseudomonas aeruginosa. Antimicrob Agents Chemother 62:1–10.

65. Ahmed MN, Porse A, Sommer MOA, Høiby N, Ciofu O. 2018. Evolution of Antibiotic Resistance in Biofilm and Planktonic Pseudomonas aeruginosa Populations Exposed to Subinhibitory Levels of Ciprofloxacin. Antimicrob Agents Chemother 62:1–12.

66. Moskowitz SM, Ernst RK, Miller SI. 2004. PmrAB, a Two-Component Regulatory System of Pseudomonas aeruginosa That Modulates Resistance to Cationic Antimicrobial Peptides and Addition of Aminoarabinose to Lipid A. J Bacteriol 186:575–579.

67. Johnson L, Mulcahy H, Kanevets U, Shi Y, Lewenza S. 2012. Surface-Localized Spermidine Protects the Pseudomonas aeruginosa Outer Membrane from Antibiotic Treatment and Oxidative Stress. J Bacteriol 194:813–826.

68. Bolard A, Schniederjans M, Haüssler S, Triponney P, Valot B, Plésiat P, Jeannot K. 2019. Production of Norspermidine Contributes to Aminoglycoside Resistance in pmrAB Mutants of Pseudomonas aeruginosa. Antimicrob Agents Chemother 63:1–14.

69. Fernández L, Gooderham WJ, Bains M, McPhee JB, Wiegand I, Hancock REW. 2010. Adaptive resistance to the “last hope” antibiotics polymyxin B and colistin in Pseudomonas aeruginosa is mediated by the novel two-component regulatory system ParR-ParS. Antimicrob Agents Chemother 54:3372–3382.

70. Puja H, Bolard A, Noguès A, Plésiat P, Jeannot K. 2020. The Efflux Pump MexXY/OprM Contributes to the Tolerance and Acquired Resistance of Pseudomonas aeruginosa to Colistin. Antimicrob Agents Chemother 64:1–11.

71. Barbosa C, Trebosc V, Kemmer C, Rosenstiel P, Beardmore R, Schulenburg H, Jansen G. 2017. Alternative Evolutionary Paths to Bacterial Antibiotic Resistance Cause Distinct Collateral Effects. Mol Biol Evol 34:2229–2244.

72. Lopatkin AJ, Bening SC, Manson AL, Stokes JM, Kohanski MA, Badran AH, Earl AM, Cheney NJ, Yang JH, Collins JJ. 2021. Clinically relevant mutations in core metabolic genes confer antibiotic resistance. Science (80-) 371.

73. Pompilio A, Crocetta V, De Nicola S, Verginelli F, Fiscarelli E, Di Bonaventura G. 2015. Cooperative pathogenicity in cystic fibrosis: Stenotrophomonas maltophilia modulates Pseudomonas aeruginosa virulence in mixed biofilm. Front Microbiol 6:1–13.

74. Magalhães AP, Lopes SP, Pereira MO. 2017. Insights into Cystic Fibrosis Polymicrobial Consortia: The Role of Species Interactions in Biofilm Development, Phenotype, and Response to In-Use Antibiotics. Front Microbiol 7:1–11.

75. Kukavica-Ibrulj I, Bragonzi A, Paroni M, Winstanley C, Sanschagrin F, O’Toole GA, Levesque RC. 2008. In Vivo Growth of Pseudomonas aeruginosa Strains PAO1 and PA14 and the Hypervirulent Strain LESB58 in a Rat Model of Chronic Lung Infection. J Bacteriol 190:2804–2813.

76. Salunkhe P, Smart CHM, Morgan JAW, Panagea S, Walshaw MJ, Hart CA, Geffers R, Tümmler B, Winstanley C. 2005. A Cystic Fibrosis Epidemic Strain of Pseudomonas aeruginosa Displays Enhanced Virulence and Antimicrobial Resistance. J Bacteriol 187:4908–4920.

77. Fothergill JL, Panagea S, Hart CA, Walshaw MJ, Pitt TL, Winstanley C. 2007. Widespread pyocyanin over-production among isolates of a cystic fibrosis epidemic strain. BMC Microbiol 7:45.

78. O’Brien S, Williams D, Fothergill JL, Paterson S, Winstanley C, Brockhurst MA. 2017. High virulence sub-populations in Pseudomonas aeruginosa long-term cystic fibrosis airway infections. BMC Microbiol 17:1–8.

79. O’Brien S, Fothergill JL. 2017. The role of multispecies social interactions in shaping Pseudomonas aeruginosa pathogenicity in the cystic fibrosis lung. FEMS Microbiol Lett 364:1–10.

80. Frimodt-Møller J, Rossi E, Haagensen JAJ, Falcone M, Molin S, Johansen HK. 2018. Mutations causing low level antibiotic resistance ensure bacterial survival in antibiotic-treated hosts. Sci Rep 8:12512.

