## Supplementary Figures and Tables for "The effects of antibiotic combination treatments on *Pseudomonas aeruginosa* tolerance evolution and coexistence with *Stenotrophomonas maltophilia*"

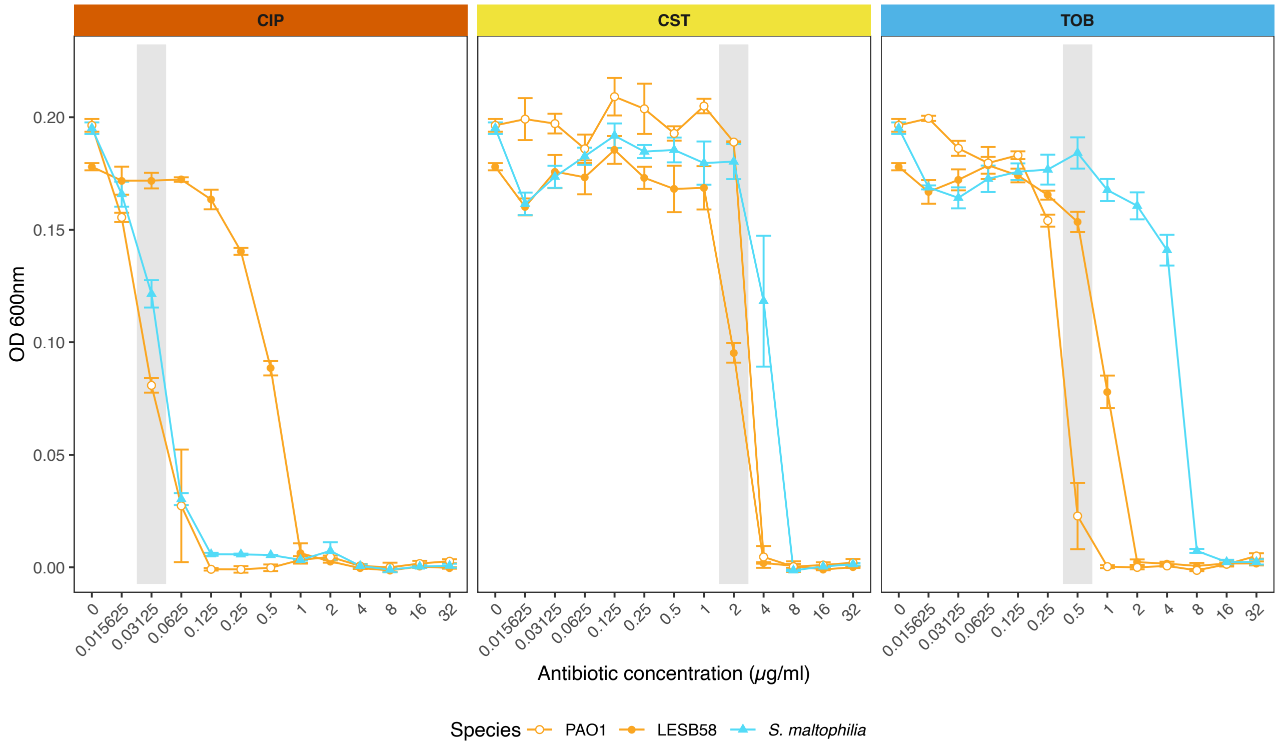


**Supplementary Figure 1: Minimum Inhibitory Concentration curves of ciprofloxacin (CIP), colistin (CST), and tobramycin (TOB) for ancestral *P. aeruginosa* strains PAO1 and LESB58, and *S. maltophilia*.**

Highlighted concentration shows the concentration used in the selection experiment. Points show means of triplicate assays, error bars ± S.E.M.

### Supplementary Tables

**Supplementary Table 1: ANOVA tables for growth of evolved *P. aeruginosa* when exposed to each antibiotic in growth assays, as difference in growth relative to ancestor. Treatment term refers to the antibiotic treatment regimen, culture refers to the presence or absence of *S. maltophilia*.**

| Strain | Environment | ANOVA term | D.F. | Sum Sq. | F value | P value |
|---|---|---|---|---|---|---|
| PAO1 | Ciprofloxacin | Treatment | 7 | 0.15 | 47.64 | $<2x10^{-16}$ |
| | | Culture | 1 | $3.9x10^{-4}$ | 0.89 | 0.35 |
| | | Interaction | 5 | $9.0x10^{-4}$ | 0.41 | 0.84 |
| | | Residuals | 48 | 0.02 | | |
| PAO1 | Colistin | Treatment | 7 | 0.06 | 6.25 | $3.15x10^{-5}$ |
| | | Culture | 1 | $9.8x10^{-5}$ | 0.08 | 0.78 |
| | | Interaction | 5 | $5.7x10^{-3}$ | 0.89 | 0.50 |
| | | Residuals | 48 | 0.06 | | |
| PAO1 | Tobramycin | Treatment | 7 | 0.18 | 15.08 | $3.14x10^{-10}$ |
| | | Culture | 1 | $1.6x10^{-3}$ | 0.99 | 0.32 |
| | | Interaction | 5 | 0.01 | 1.38 | 0.25 |
| | | Residuals | 48 | 0.08 | | |
| PAO1 | No Antibiotic | Treatment | 7 | $7.7x10^{-3}$ | 5.06 | $2.3x10^{-4}$ |
| | | Culture | 1 | $1.1x10^{-6}$ | $5.0x10^{-3}$ | 0.94 |
| | | Interaction | 5 | $5.7x10^{-4}$ | 0.52 | 0.76 |
| | | Residuals | 48 | 0.01 | | |
| LESB58 | Ciprofloxacin | Treatment | 7 | 0.16 | 7.25 | $1.96x10^{-6}$ |
| | | Culture | 1 | $3.8x10^{-5}$ | 0.01 | 0.91 |
| | | Interaction | 7 | $3.9x10^{-3}$ | 0.17 | 0.99 |
| | | Residuals | 66 | 0.21 | | |
| LESB58 | Colistin | Treatment | 7 | 0.16 | 8.14 | $4.08x10^{-7}$ |
| | | Culture | 1 | $3.1x10^{-5}$ | 0.01 | 0.92 |
| | | Interaction | 7 | 0.02 | 0.81 | 0.59 |
| | | Residuals | 66 | 0.19 | | |
| LESB58 | Tobramycin | Treatment | 7 | 0.31 | 14.75 | $2.07x10^{-11}$ |
| | | Culture | 1 | $5.9x10^{-4}$ | 0.20 | 0.66 |
| | | Interaction | 7 | $6.2x10^{-3}$ | 0.30 | 0.95 |

| | | | | | | |
|---|---|---|---|---|---|---|
| | | Residuals | 66 | 0.20 | | |
| LESB58 | No Antibiotic | Treatment | 7 | 0.02 | 6.25 | $1.24x10^{-5}$ |
| | | Culture | 1 | $2.5x10^{-5}$ | 0.06 | 0.81 |
| | | Interaction | 7 | $3.9x10^{-3}$ | 1.39 | 0.23 |
| | | Residuals | 66 | 0.03 | | |

**Supplementary Table 2: Pearson Chi-Squared Test of Independence for the MIC of each antibiotic for both *P. aeruginosa* strains.**

| Strain | Antibiotic | $X^2$ | D.F. | P value |
|---|---|---|---|---|
| PAO1 | Ciprofloxacin | 32.13 | 7 | $3.84x10^{-5}$ |
| | Colistin | 29.00 | 7 | $1.44x10^{-4}$ |
| | Tobramycin | 30.55 | 7 | $7.52x10^{-5}$ |
| LESB58 | Ciprofloxacin | 33.47 | 7 | $2.16x10^{-5}$ |
| | Colistin | 29.89 | 7 | $9.97x10^{-5}$ |
| | Tobramycin | 37.86 | 7 | $3.22x10^{-6}$ |

**Supplementary Table 3 ANOVA tables of natural logarithm transformed total population density (as $OD_{600}$) at the final timepoint of the selection experiment. Treatment term refers to the antibiotic treatment regimen, culture refers to the presence or absence of *S. maltophilia*.**

| Strain | ANOVA term | D.F. | Sum Sq. | F value | P value |
|---|---|---|---|---|---|
| PAO1 | Treatment | 7 | 1.67 | 4.42 | 0.00045 |
| | Culture | 1 | 0.0024 | 0.044 | 0.83 |
| | Interaction | 7 | 0.22 | 0.58 | 0.77 |
| | Residuals | 65 | 3.50 | | |
| LESB58 | Treatment | 7 | 2.81 | 8.30 | $2.29x10^{-7}$ |
| | Culture | 1 | 0.28 | 5.51 | 0.022 |
| | Interaction | 7 | 0.53 | 1.57 | 0.16 |
| | Residuals | 71 | 3.44 | | |
